## Supplementary figures and images for "Computational design of novel Cas9 PAM-interacting domains using evolution-based modelling and structural quality assessment"

### dg_1_plddt.png

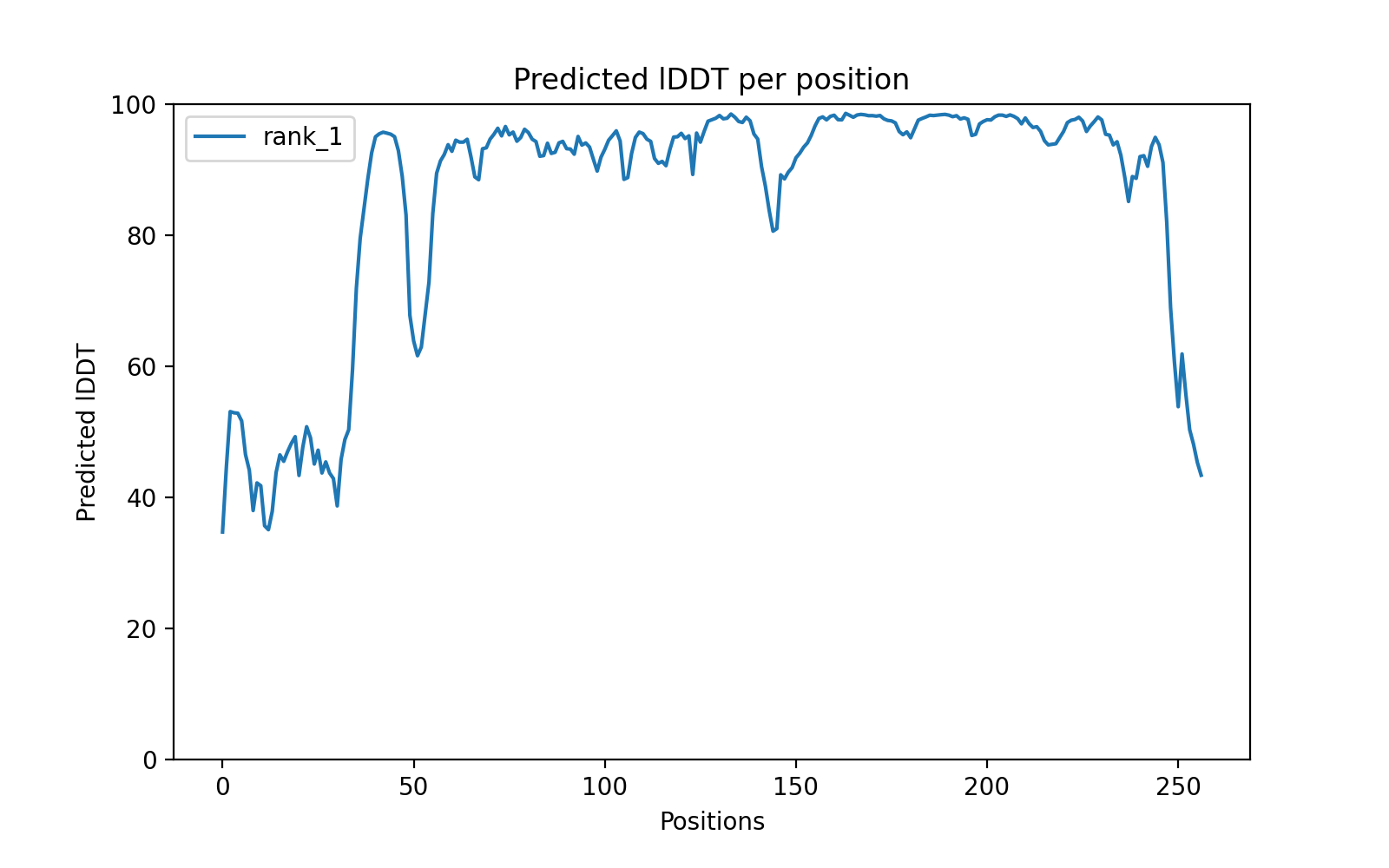

### dg_3_plddt.png

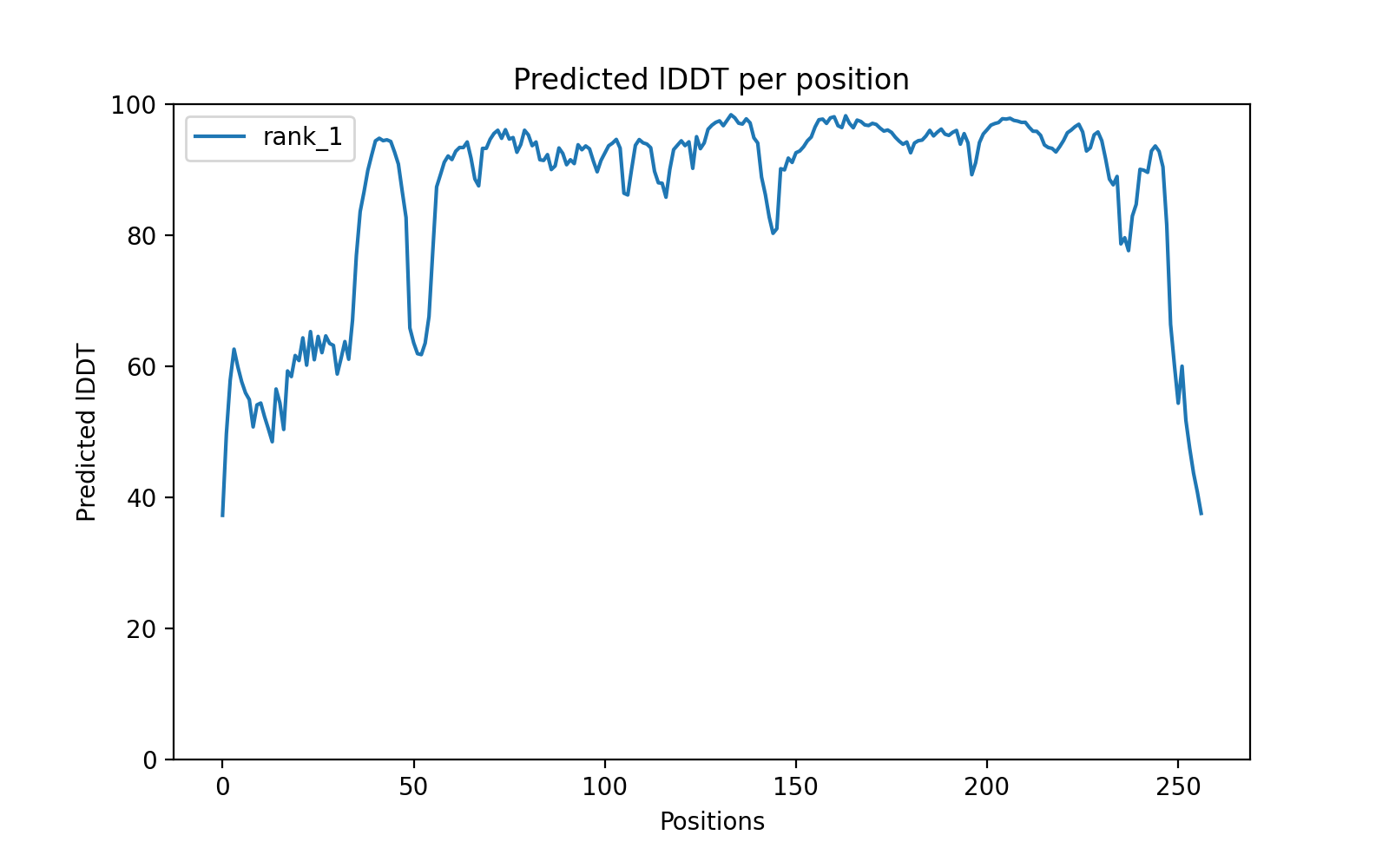

### dg_9_plddt.png

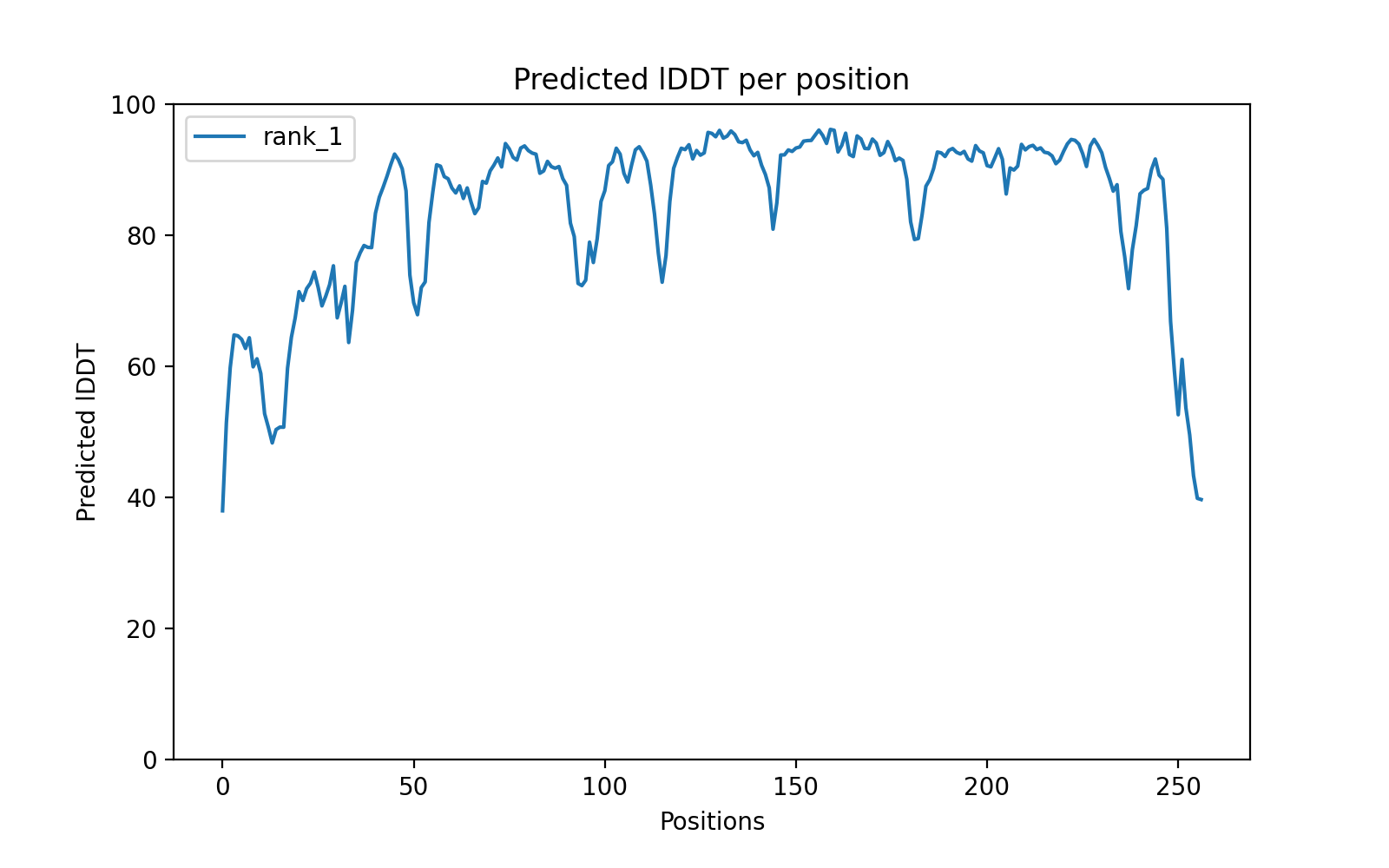

### dg_10_plddt.png

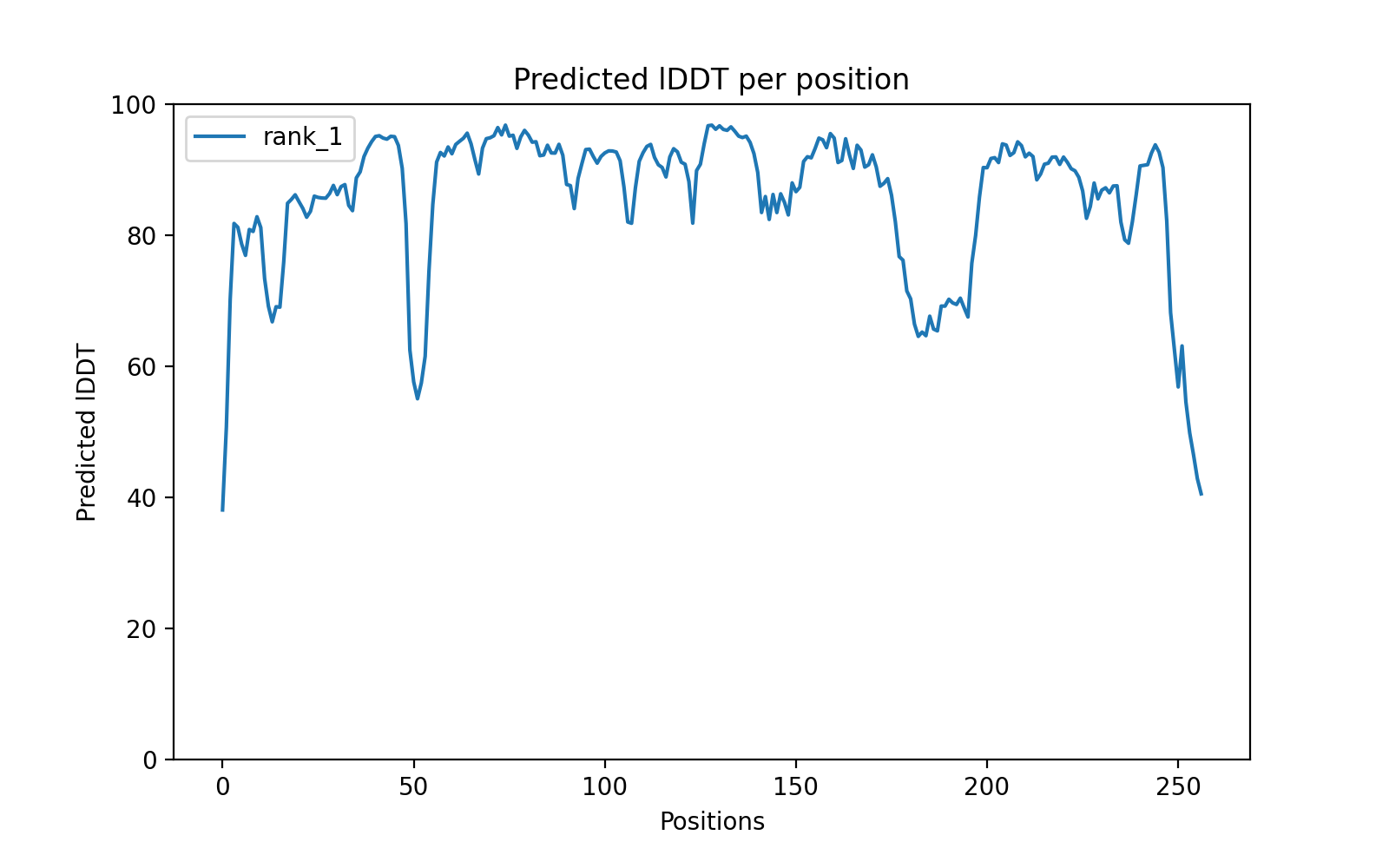

### dg_15_plddt.png

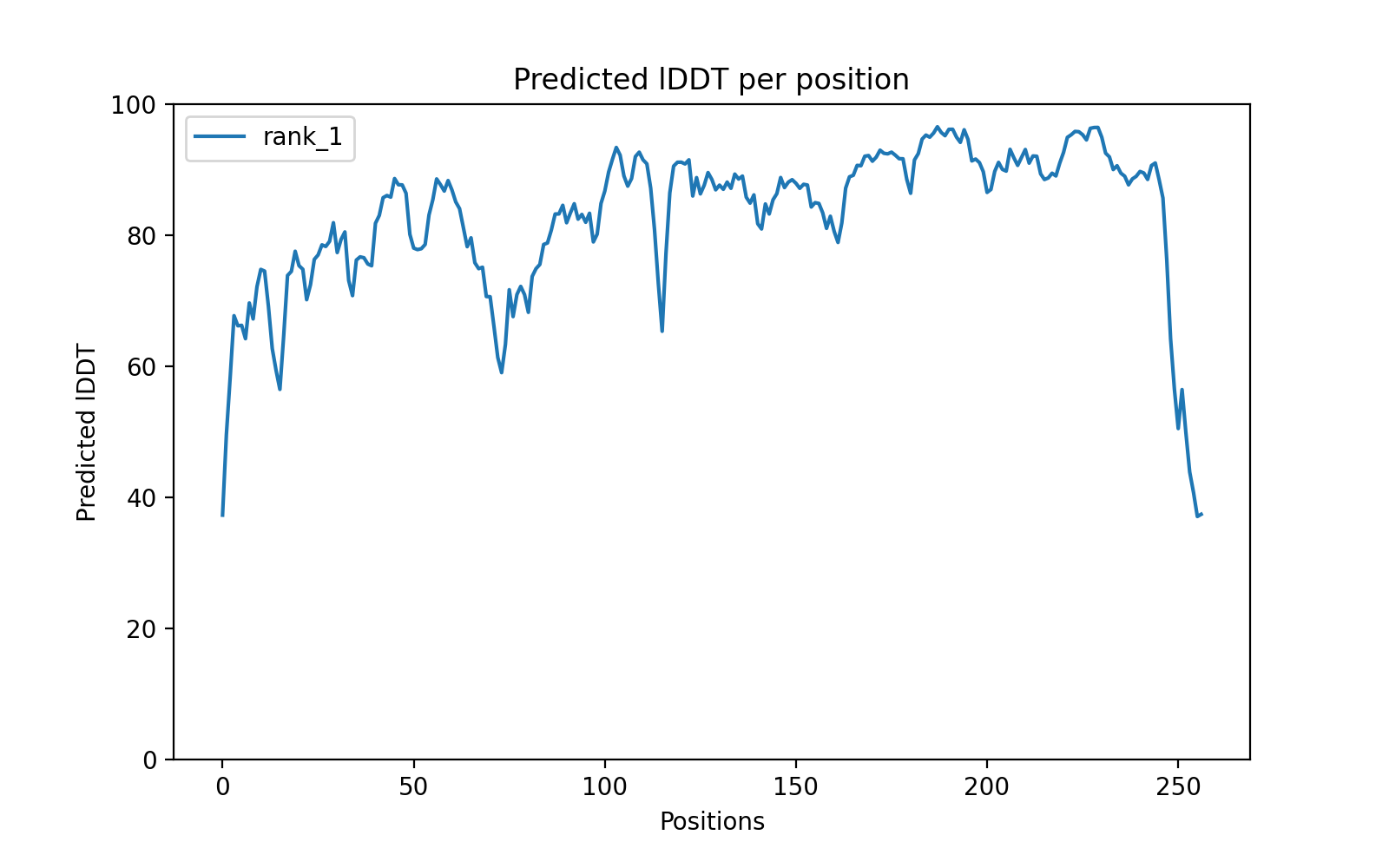

### dg_18_plddt.png

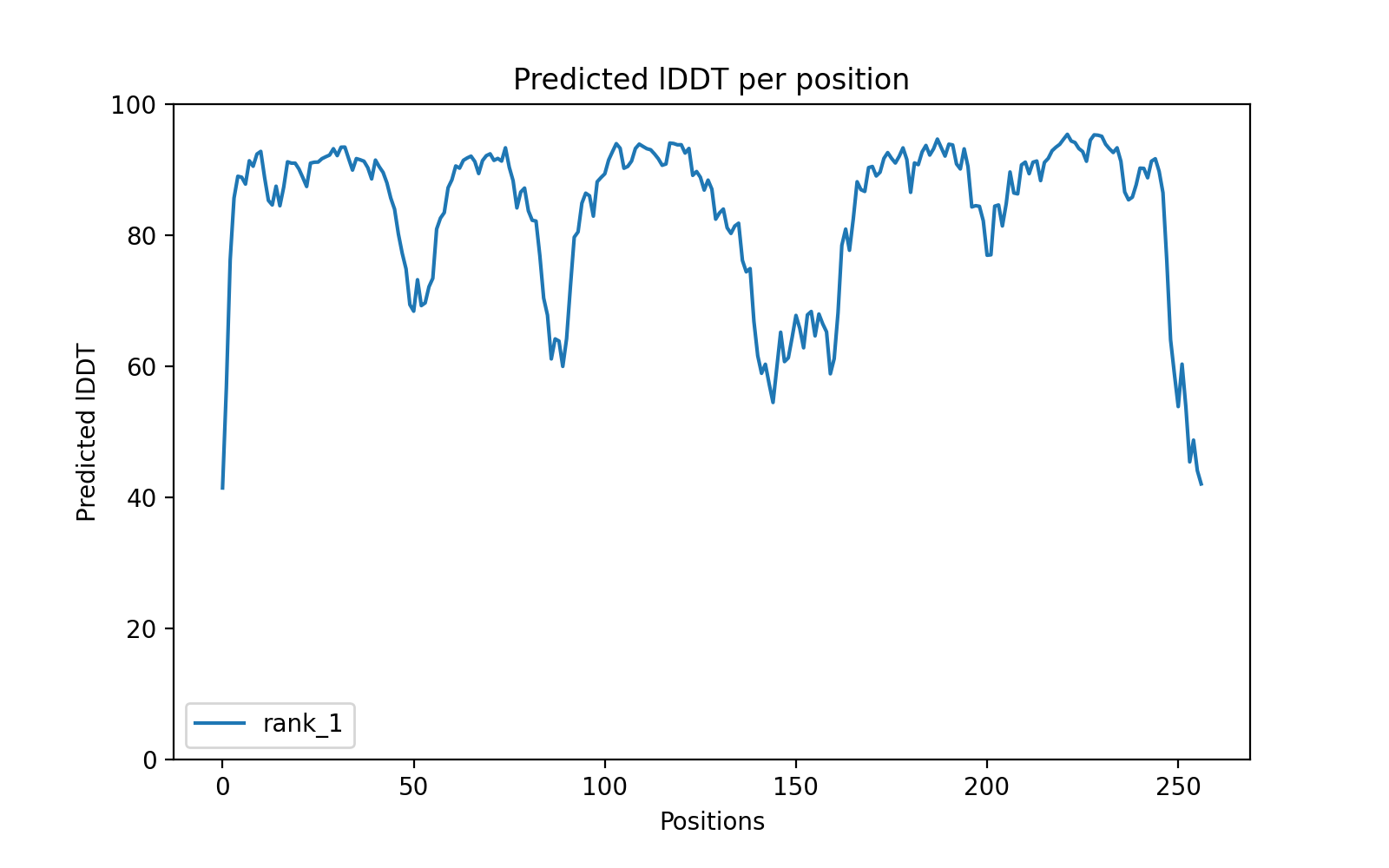

### dg_22_plddt.png

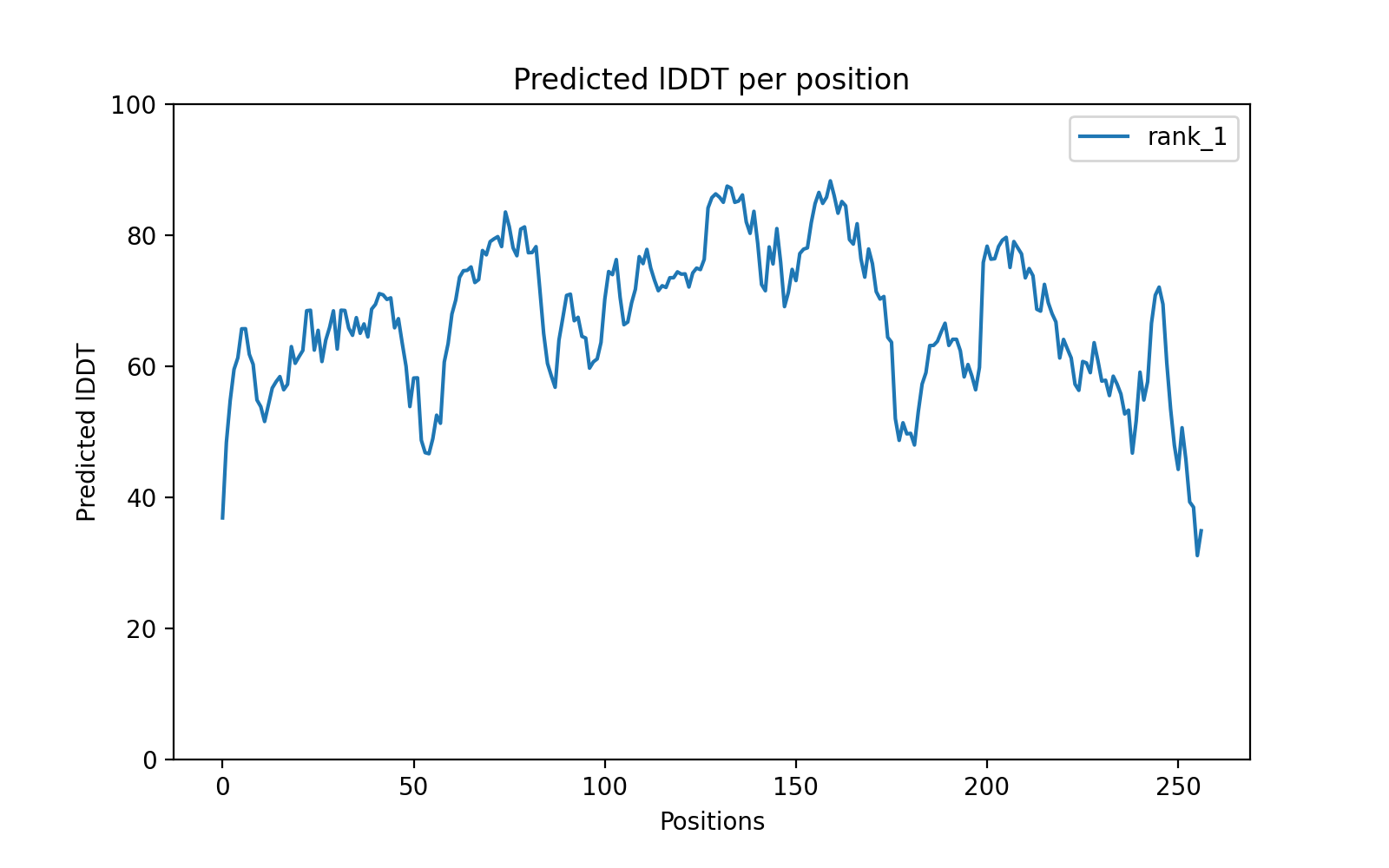

### dgfx_7_plddt.png

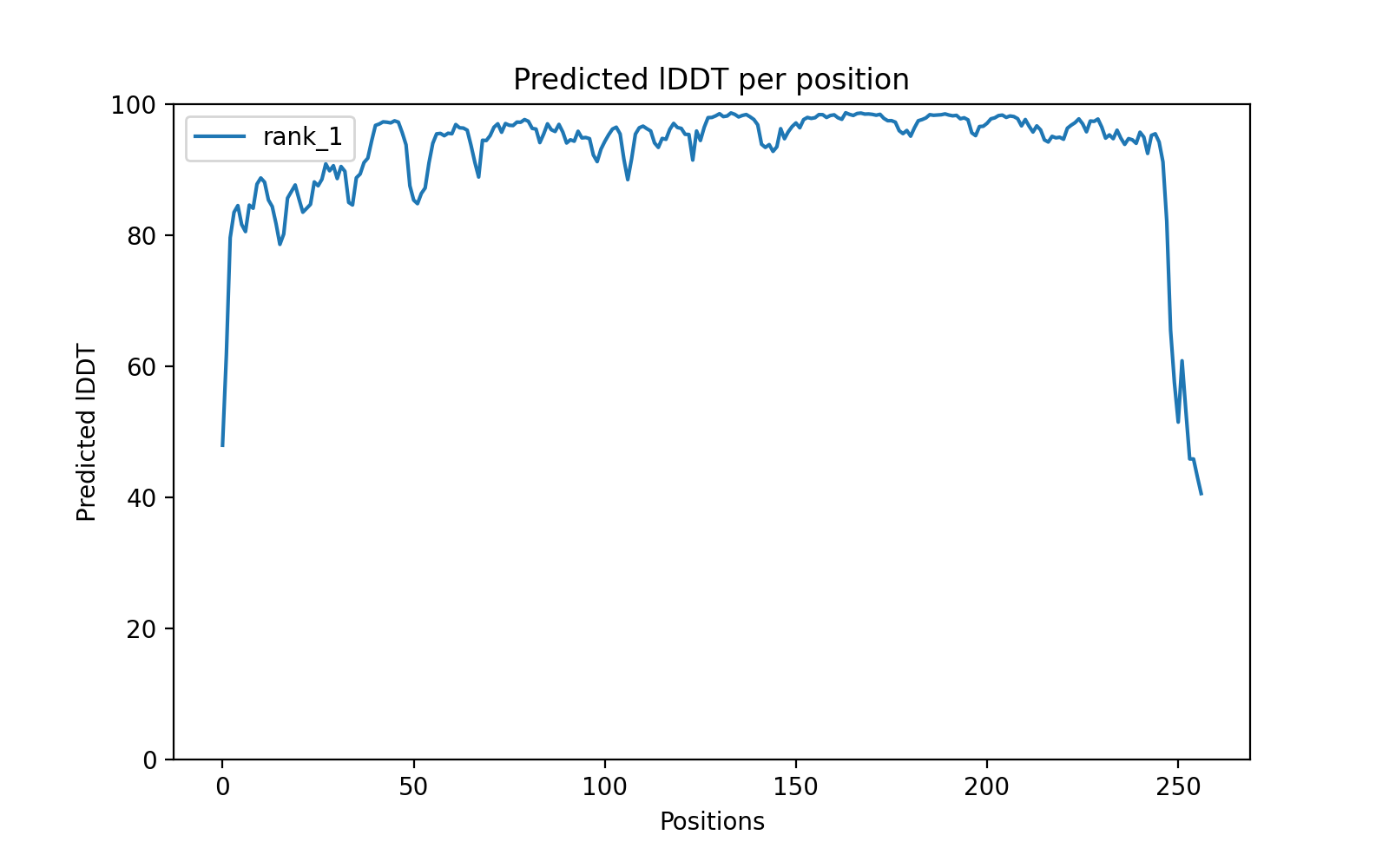

### dgfx_15_plddt.png

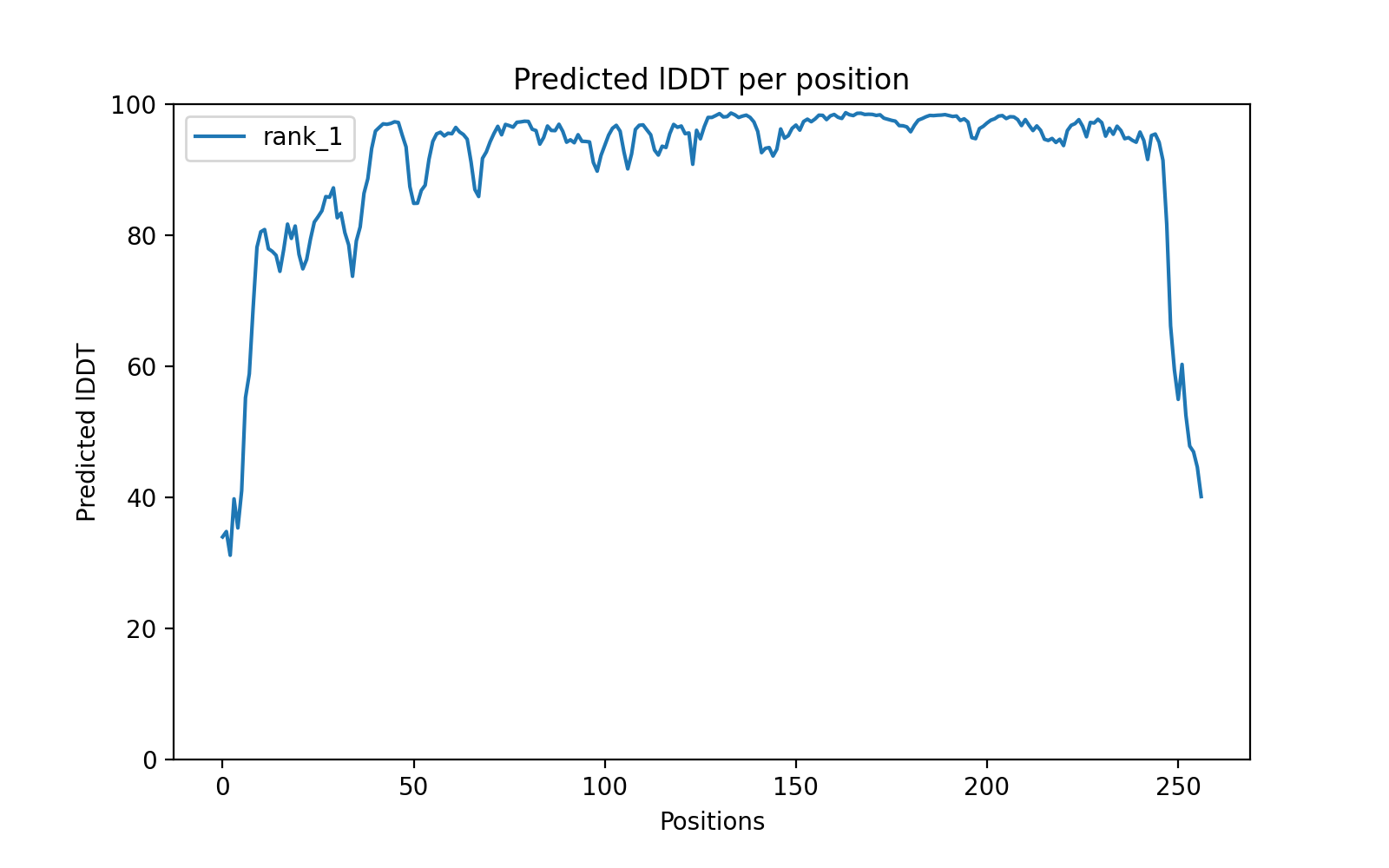

### dgfx_22_plddt.png

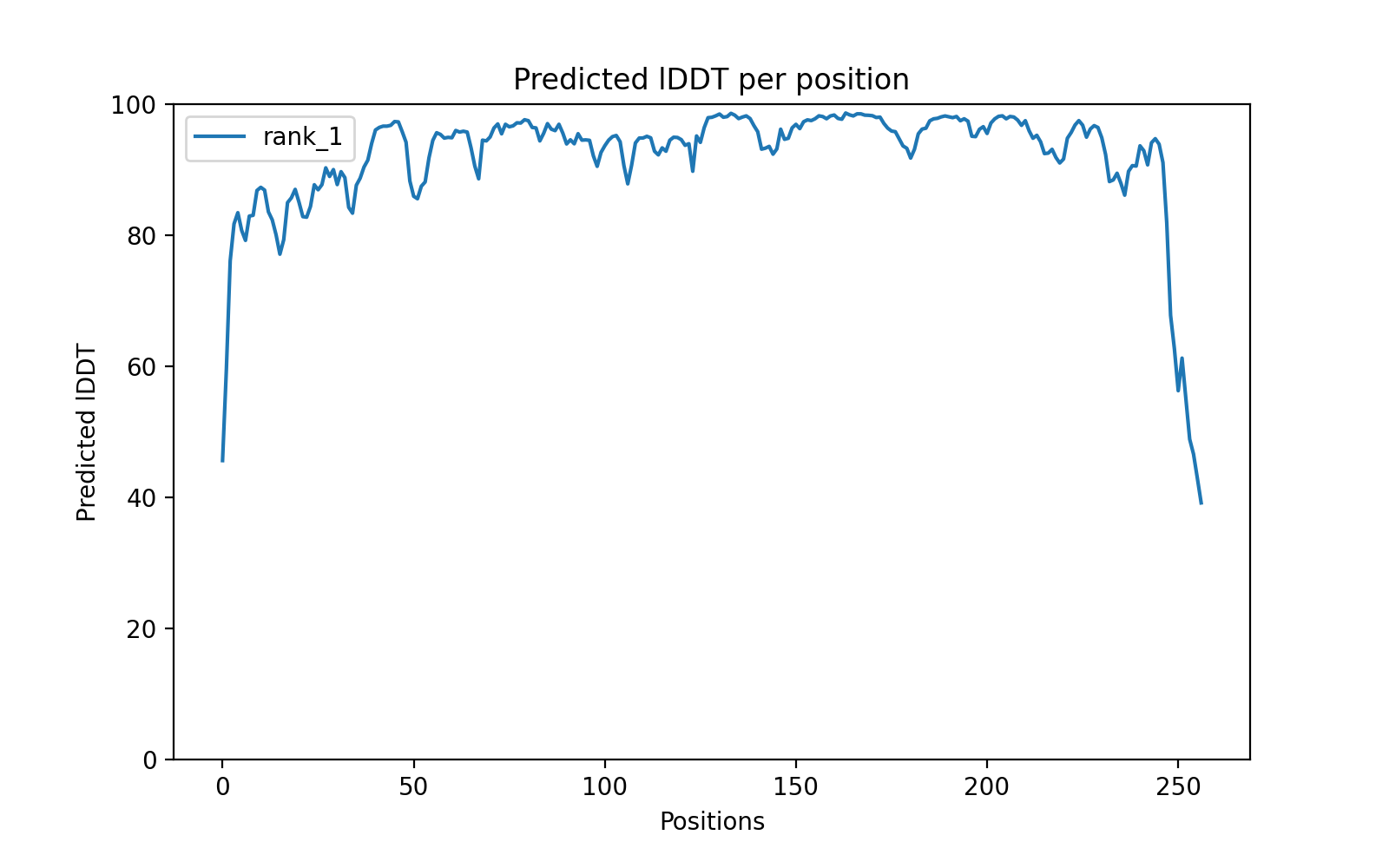

### dgfx_34_plddt.png

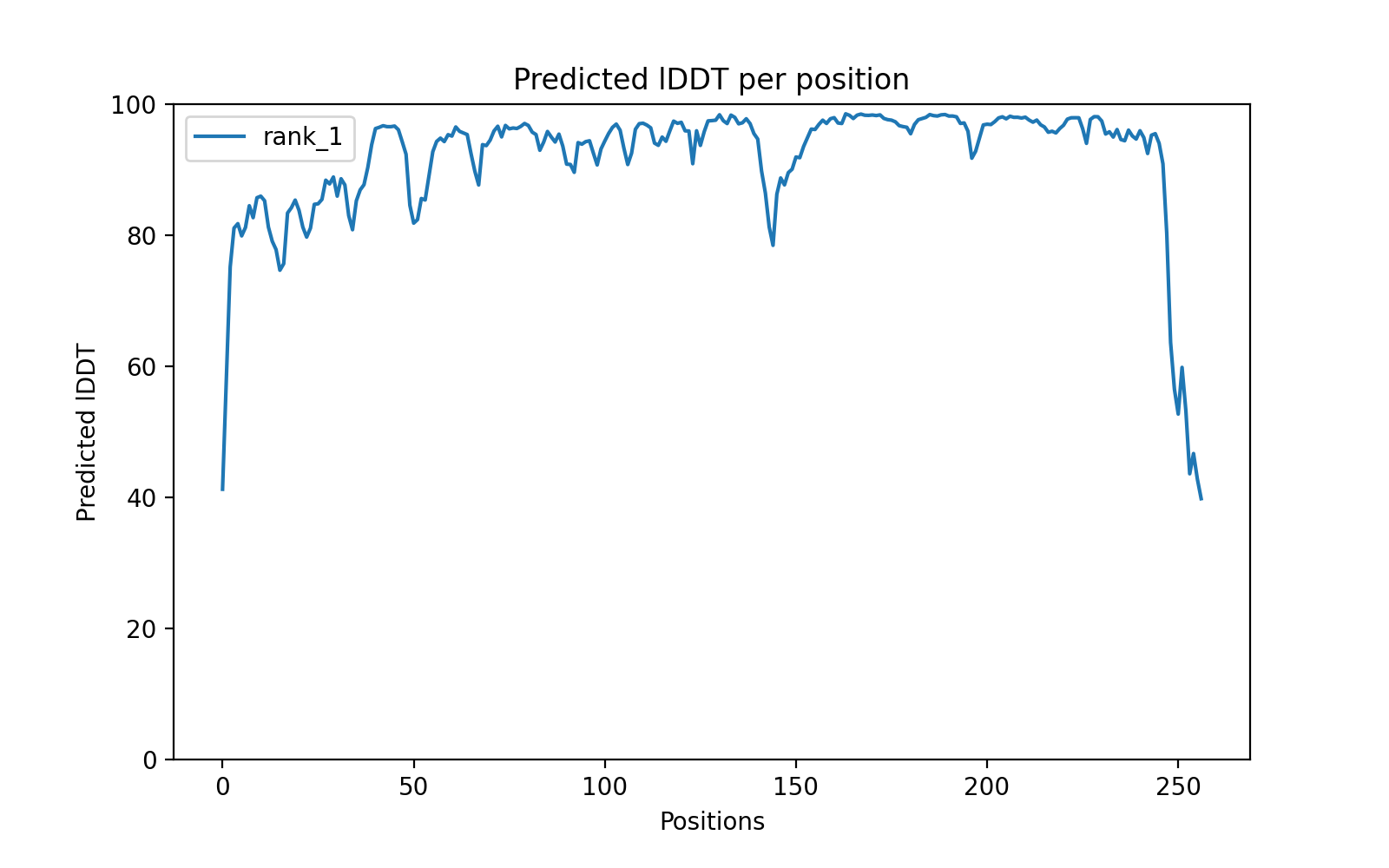

### dgfx_69_plddt.png

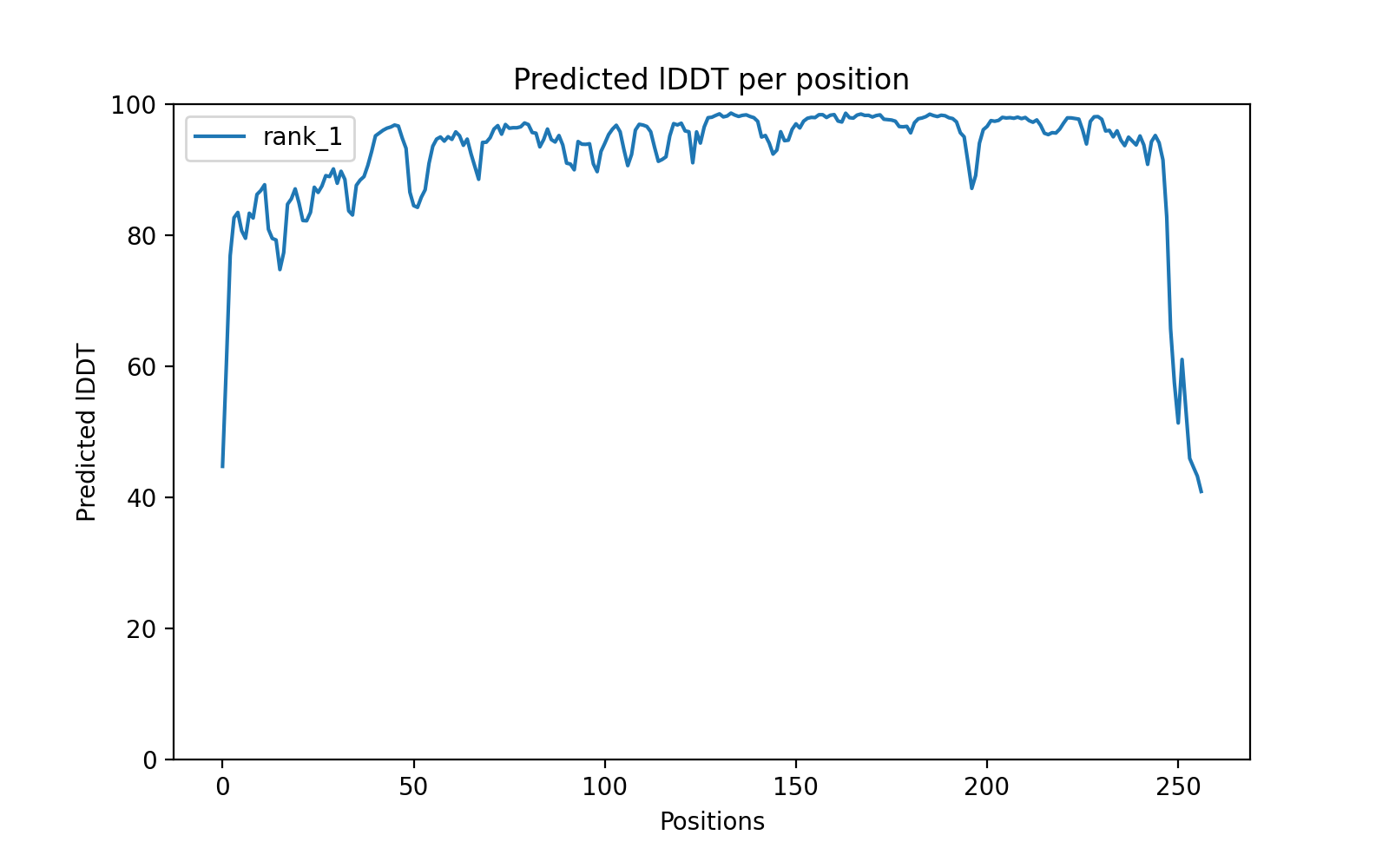

### S6 file

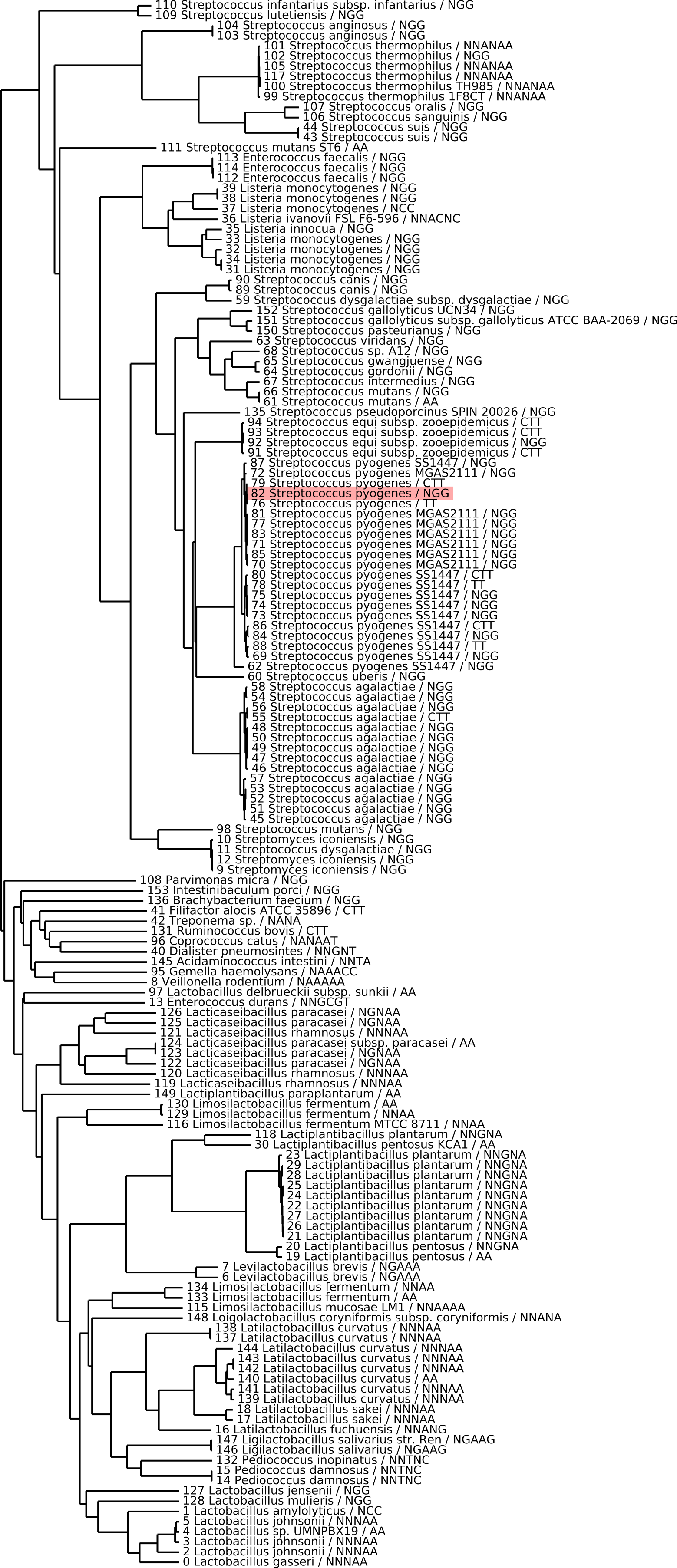
