## Supplementary material for "Computational design of novel Cas9 PAM-interacting domains using evolution-based modelling and structural quality assessment": S3 file

### S3 file : Additional plots on assessments of computational score

### 1 Prediction of activity using different scores

We performed logistic regression using different combinations of scores.We first used the first batch as a training set and the second batch as a testing set to demonstrate that the results of the second batch were largely predictable from those of the first batch. We also performed a completely randomized 2-fold cross validation (reproduced 100 times)

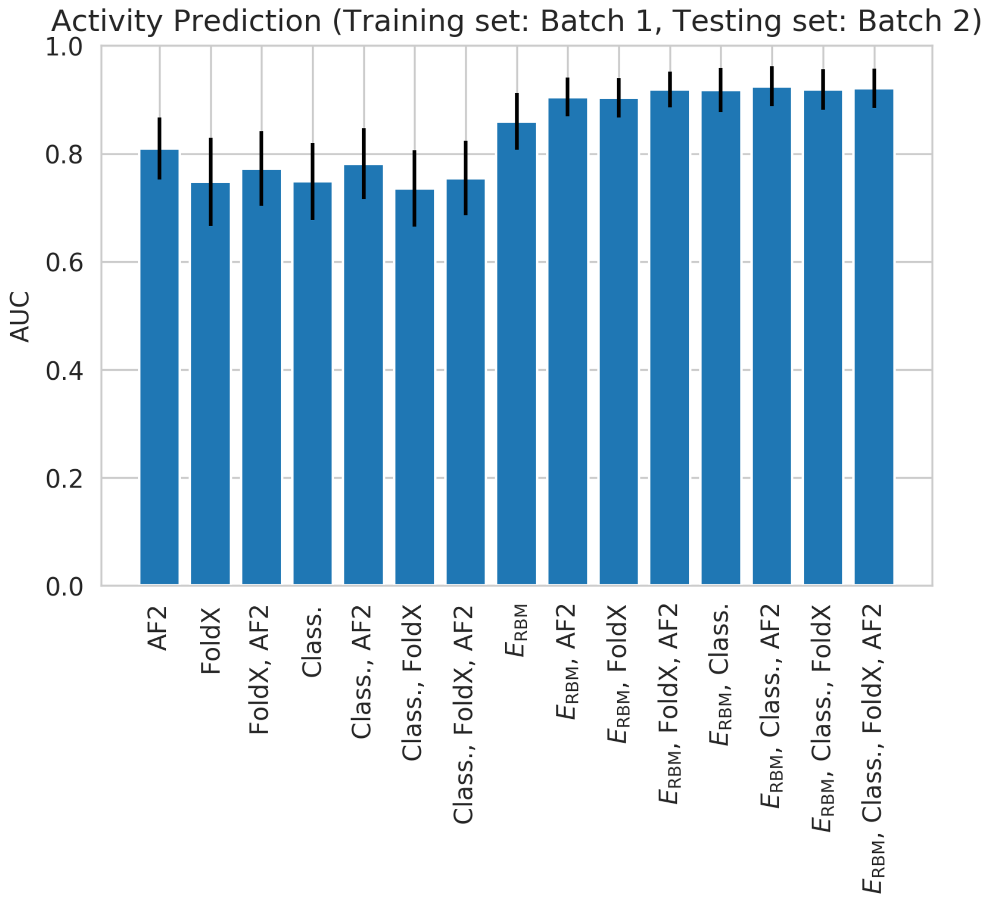


**Fig 1. a**: Activity prediction ($\mathrm{mcherry} \geq 0.5$) trained using logistic regression. The training set is the first batch, the testing set is the second batch. **b**: Activity prediction ($\mathrm{mcherry} \geq 0.5$) trained and tested through cross validation (completely random, 2 fold, reproduced 100 times).

#### 2 Prediction of activity through different scores

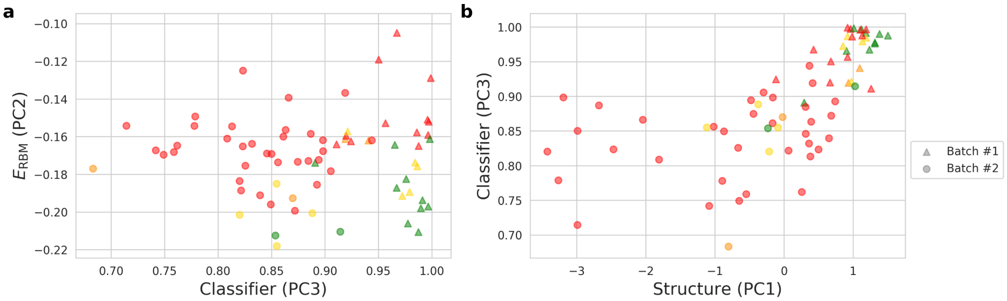


**Fig 2. a**: The classifier score and the RBM energy are not correlated and carry complementary information for activity prediction. **b**: The classifier score correlates with the structural scores.
.
