## Supplementary material for "Computational design of novel Cas9 PAM-interacting domains using evolution-based modelling and structural quality assessment": S4 file

### S4 file : Experimental measurements of the activity through both mCherry and sacB

#### 1 Plasmid used for experimental validation

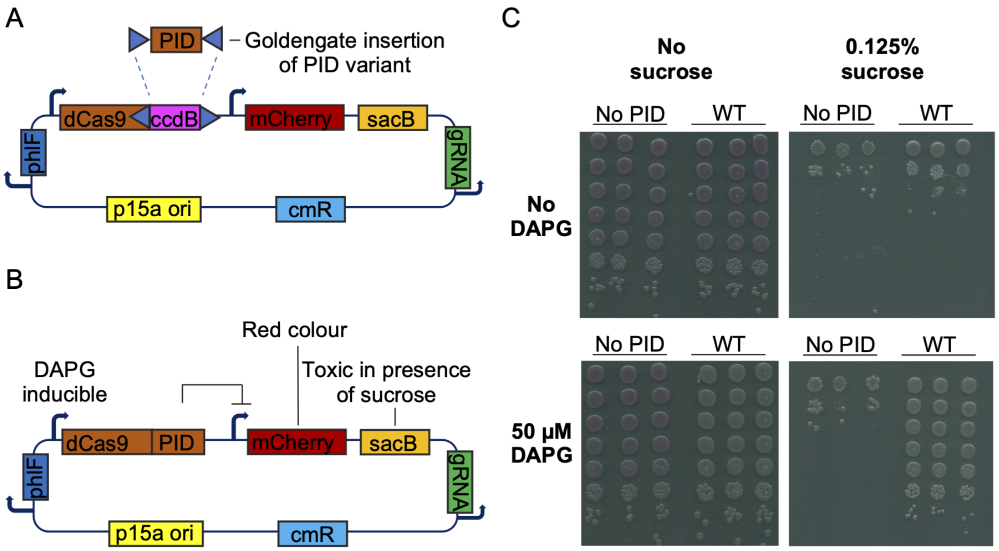


**Fig 1. A**: Diagram of pWR5, which contains a goldengate site for easy cloning and testing of PID fusions. A guide RNA targets the mCherry-sacB promoter. **B**: Circuit after cloning of a PID variant. **C**: Spot assay of either pWR8 (containing the WT *S. pyogenes* Cas9) or the inactive control pWR9, where GFP is cloned instead of an active PID, on plates with or without 50 $\mu$M DAPG and with or without 0.125% sucrose

.

#### 2 Trees of natural Cas9 PID tested as chimera with SpyCas9

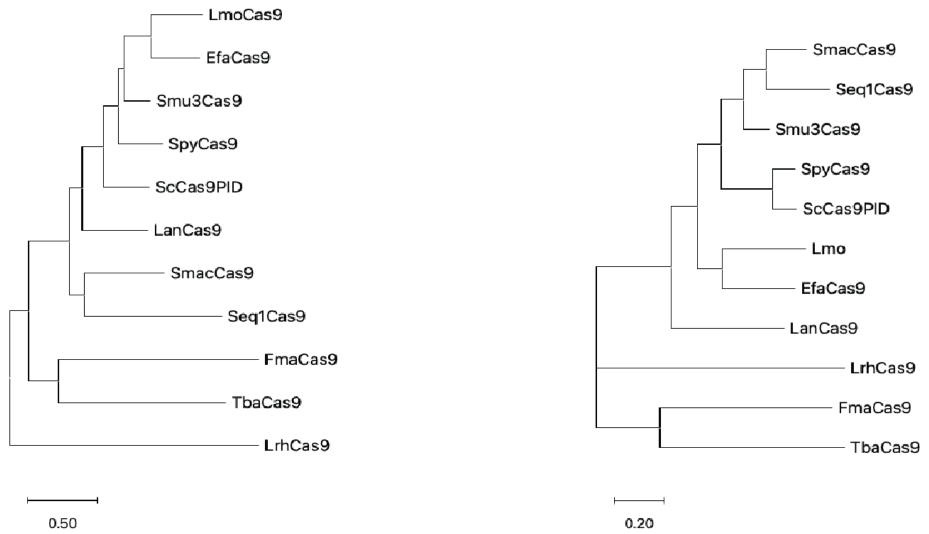


**Fig 2.** Maximum-likelihood trees of tested natural PID chimeras, for either whole Cas9 proteins (left) or only the PAM-interacting domains (right). Maximum-likelihood phylogenetic trees were built using MEGA11.

#### 3 Pictures of tested enzymes

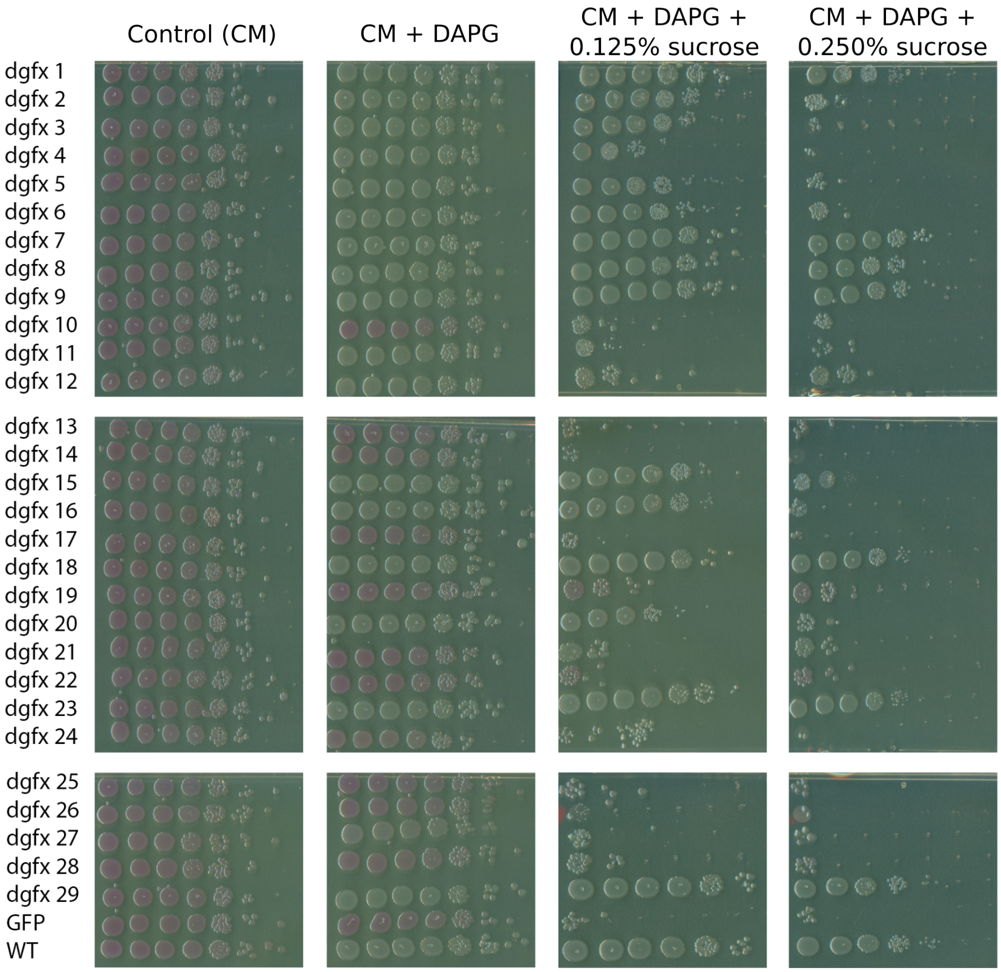


**Fig 3.** Experimental result of the first batch. Generated sequences were cloned in pWR5 and introduced in E. coli MG1655. Overnight cultures were serially diluted (10x dilutions at each step) and spotted on plates supplemented with DAPG (to induce Cas9 expression) and sucrose as indicated.

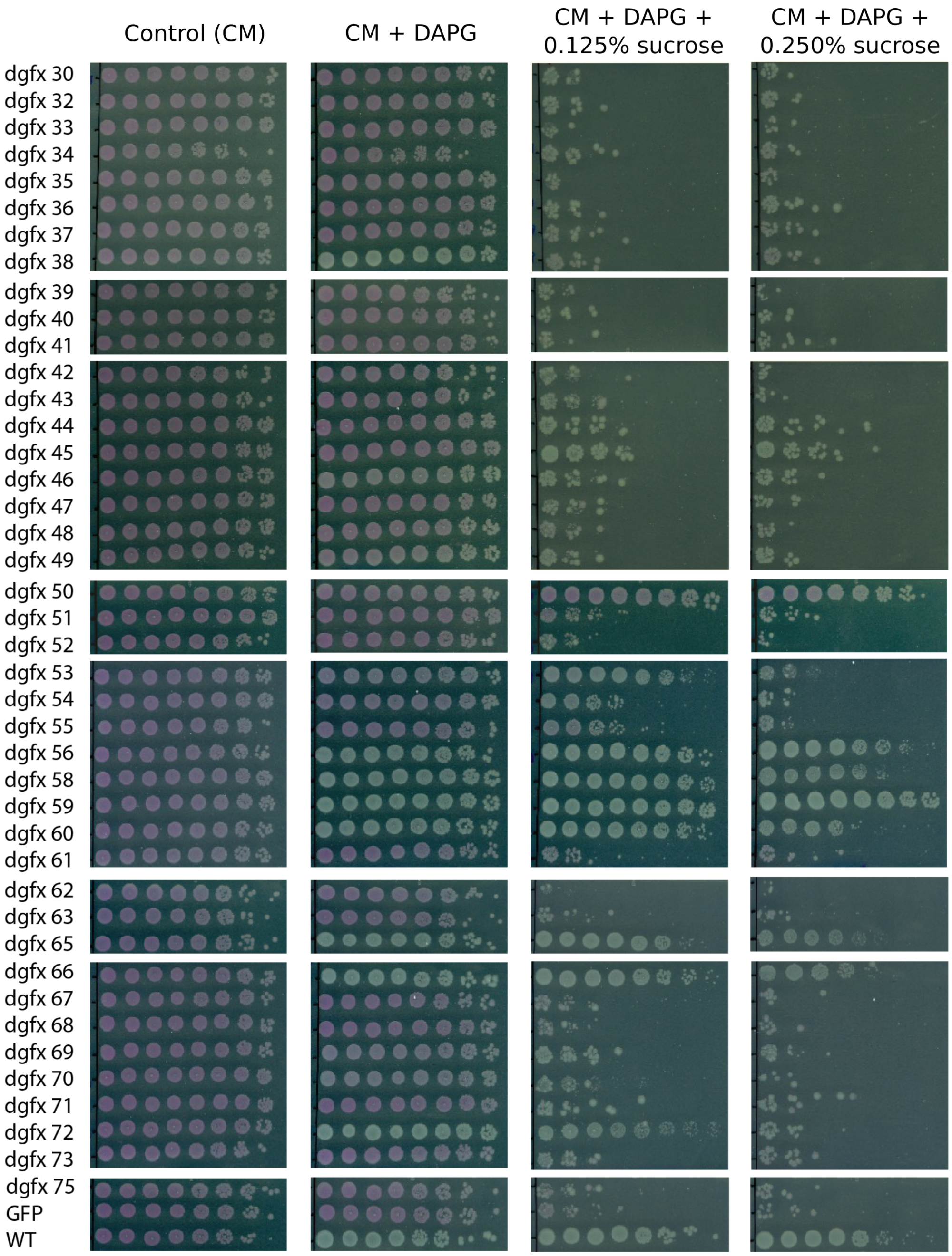


**Fig 4.** Experimental result of the second batch. Generated sequences were cloned in pWR5 and introduced in E. coli MG1655. Overnight cultures were serially diluted (10x dilutions at each step) and spotted on plates supplemented with DAPG (to induce Cas9 expression) and sucrose as indicated.
