## Supplementary material for "Computational design of novel Cas9 PAM-interacting domains using evolution-based modelling and structural quality assessment": S5 file

### S5 file : application to MNIST, towards building transition path

To illustrate the ability of our Constrained Langevin Dynamics to follow complex constraints, we propose to see how it can solve the difficult problem of finding a transition path between two almost disconnected modes of a distribution.

We take an SSL-RBM trained on zeros and ones digits of MNIST, and we look for a transition path between a specific 0, $x^{(0)}$, and a specific 1, $x^{(1)}$, which includes intermediate samples with high likelihoods according to the RBM. To do so, we resort to the following criteria.

#### 1 Method 1: Use distance to $x^{(1)}$ as a guide

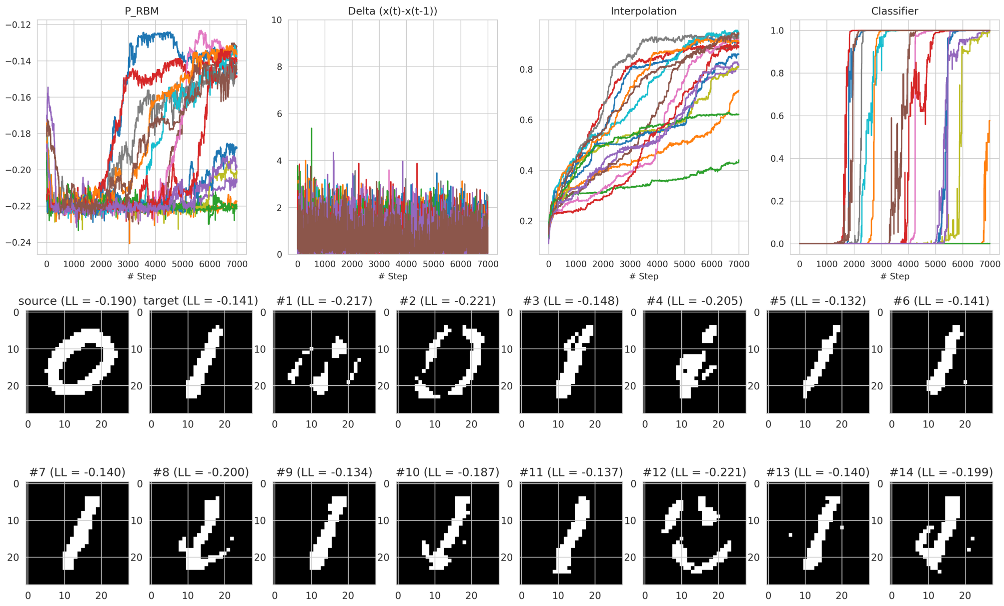


**Fig 1.** We display some transition paths from Constrained Langevin Dynamics using method 1. Many chains stay stuck and do not find a transition path, but some succeed to reach the target while respecting specific conditions (high likelihood samples + small changes at each step)

Our main criterion guides the chain from $x^{(0)}$ to $x^{(1)}$:

- The transition ratio: we define the transition ratio by the function $f_1$:

$$f_1(x) = \frac{\sum_i \mathbf{1}(x_i \neq x_i^{(0)})}{\sum_i \mathbf{1}(x_i \neq x_i^{(0)}) + \sum_i \mathbf{1}(x_i \neq x_i^{(1)})} \tag{1}$$

  This function is for example null for $x = x^{(0)}$, equal to one for $x = x^{(1)}$ and if $x = 0.5$, $x$ is equally distant to both.

The next set of criteria are those we want to control (by convention, their values are set to zero):

- The delta criterion: we want the transition path to change a few pixels (say, less than $\delta_{max} = 5$) at each step. Let $x^{(t)}$ be the element of the chain at time $t$, we define the delta criterion to be:

$$\delta_t(x, x^{(t)}) = (\sum_i \mathbf{1}(x_i^{(t)} \neq x_i) - \delta_{max})^+ \tag{2}$$

  This function will be null if the number of change between two consecutive states of the chain has less than five pixels of difference.

- The RBM likelihood criterion: we want the RBM likelihood of our samples to be above a certain threshold $p_{min}$. This means that we want to keep null following criterion:

$$p(x) = (p_{min} - \mathbb{P}_{RBM}(x))^+ \tag{3}$$

In Figure 1, we display the result of this method. Overall, a fair share of chains were able to perform a transition between the source and the target and the control criterion ($p_{min} = -0.22$, $\delta_{max} = 5$) were mostly respected.

#### 2 Method 2: Use classifier as a guide

Another method allows us to have a "linear" transition path between $x^{(0)}$ to $x^{(1)}$. Instead of using the transition ratio as the main criterion, we consider as one of our dynamical criteria to control. Our main criterion is then the classifier use to direct us in the direction of the 1. While the criteria to control are now three:

- The delta criterion: unchanged
- The RBM likelihood criterion: unchanged
- The transition ratio: we want the transition to happen linearly in $N$ steps, we then make so that at each time $t$ the transition ratio be around the ratio of the number of steps it is expected to take.

$$f_t(x) = \left( \frac{\sum_i \mathbf{1}(x_i \neq x_i^{(0)})}{\sum_i \mathbf{1}(x_i \neq x_i^{(0)}) + \sum_i \mathbf{1}(x_i \neq x_i^{(1)})} - t/N \right)^2 \tag{4}$$

We see in Figure 2 that this method sometimes fails to meet some criteria, but discrepancies are still reasonable. The transition, however, is way more controlled, with all the chains able to follow the transition path at the expected rhythm.

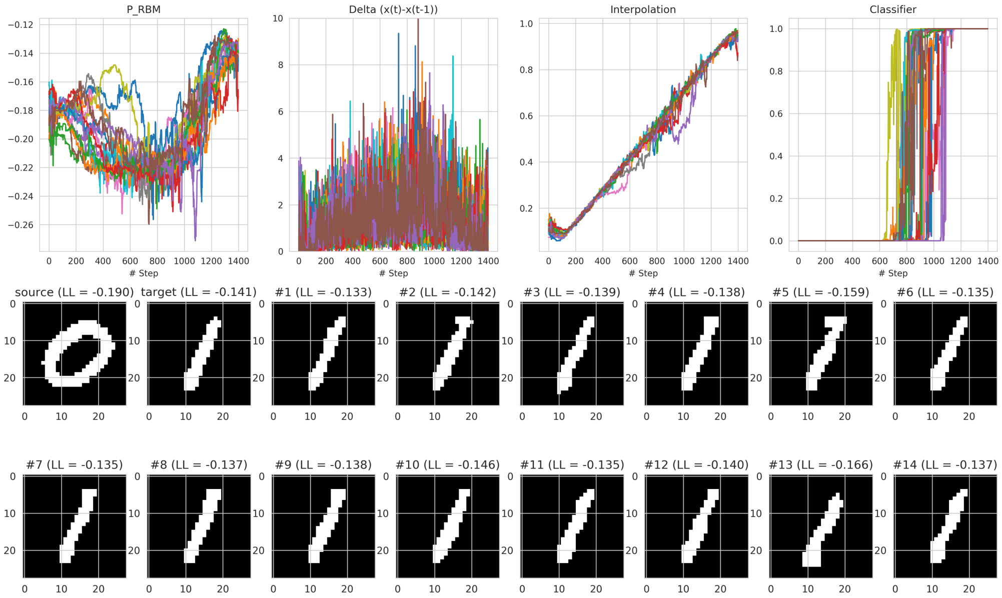


**Fig 2.** We display some transition paths built with Constrained Langevin Dynamics using method 2. The chains are really respectful of the transition ratio, but occasionally, they broke the constraints on the likelihood of the samples ($p_{min} = -0.22$) or on the small changes ($\delta_{max} = 5$).
